# Changes in plant physiology during cultivation periods influence the hydrogen isotope ratio of the water-insect relationship

**DOI:** 10.1101/2024.03.06.583809

**Authors:** Tomohisa Fujii, Gaku Akiduki, Shiho Yabusaki, Ichiro Tayasu

## Abstract

1. The hydrogen stable isotope ratios (δ^2^H) in tissues of terrestrial insects are widely applied to estimate natal origins in field populations. The hydrogen isotopes of insect tissues incorporate those of environmental waters through the insects’ metabolic processes. Water sources and abiotic environmental factors reflect changes in plant physiology, as indicated by the δ^2^H values of plants. However, the influence of plant physiology on the assimilation of hydrogen in insect tissues derived from water through feeding diets is still unknown.
2. We experimentally examined the influence of water on the δ^2^H values of maize leaf and of forewings of *Mythimna separata* (Lepidoptera: Noctuidae) and *Spodoptera frugiperda* (Noctuidae). We prepared five specific water samples for cultivating maize, aiming to replicate the gradient of δ^2^H values observed in environmental waters across the Japanese archipelago.
3. The mean-percentage contribution of water to hydrogen in maize leaves was 17.6% (July-August) and 25.1% (September-October). Linear analyses indicated that 17.4% and 32.7% of hydrogen in *M. separata* and *S. frugiperda* forewings were derived from water through the consumption of maize leaves. The slope values of linear regression between the insect forewings and the maize leaves supplied in the final instar were closest to 1.0. These results indicated that the δ^2^H values of maize leaves and insect forewings reflected those of water resources.
4. This study presented the seasonal changes in climate conditions that affected the δ^2^H values of host plants and insect tissues. Changes in plant physiology with seasonal variations may influence the interpretation of the linear relationship between water and insect tissues on the estimation of natal origins.

## Introduction

The hydrogen stable isotope is a good tool for tracing the seasonal migration of birds and insects between continents (Hobson & Wassenaar, 2018). The hydrogen isotopes derived from water in plants and animal tissues are assimilated through water‒ plant‒animal relationships. The hydrogen stable isotope ratios (δ^2^H) of plants usually reflect those of environmental waters because xylem water is typically unfractionated relative to the soil water and groundwater (Bowen, 2010). A map illustrating the δ^2^H values of water sources along environmental gradients reveals spatial patterns and spatiotemporal variability in the preferential utilization of water sources and their contribution to plant growth (Bowen, 2010). The δ^2^H values of animal tissues (e.g., animal bone, bird feather, beetle and butterfly wings) that are correlated with those of the environmental waters are analogous to the δ^2^H values of water‒plant relationships (e.g., Hobson et al., 1999a, b; 2018; Gröcke et al., 2006; Clem et al., 2023). Such maps can therefore be used to infer the locations of natal origin in animal migration studies (Hobson & Wassenaar, 2018).

For migration studies, the hydrogen stable isotopes in metabolically inert animal tissues (e.g., enamel, keratin, chitin) assume the incorporation of hydrogen derived from local diets as the isotopic signatures of the natal origin of animal growth (Hobson & Wassenaar, 2018). The basis of this assumption is that the role of δ^2^H values of water in animal tissues has been investigated by the interpretation of slope values of linear model analyses between the δ^2^H values of water and animal tissues in various species for three decades (Hobson et al., 1990a; Ehleringer et al., 2008; Wolf et al., 2011). Experimental studies showed that 30% to 50% of hydrogen was incorporated from water sources through the consumption of water or food in birds (Hobson et al., 1999a; Wolf et al., 2011), humans (Ehleringer et al., 2008), aquatic invertebrates (Nielson & Bowen, 2010; Solomon et al. 2009; Wang et al., 2009), and terrestrial insects (Hobson et al.,1999b, 2018; Holder, 2012; Clem et al., 2022). To understand the natal origin of animal growth, the isotopic composition of hydrogen derived from water in diets and animal tissues must be revealed.

The long-distance migration of *Mythimna separata* (Walker) (Lepidoptera: Noctuidae) and *Spodoptera frugiperda* (J. E. Smith) (Noctuidae) is a commonly observed phenomenon (e.g., Koyama & Matsumura, 2019; Gao et al., 2020). *Mythimna separata*, the oriental armyworm, is distributed in South, Southeast, and East Asia; New Guinea, Australia, and New Zealand; and the Pacific islands (Koyama & Matsumura, 2019). *Spodoptera frugiperda*, the fall armyworm, is distributed across a wide area of South, middle-South, and North America, as well as in Africa (Tay et al., 2023). Since 2018, *S. frugiperda* has invaded from eastern Africa to India, Myanmar, China, Vietnam, South Korea, Japan, Indonesia, Australia, and New Zealand (see the summary in Tay et al., 2022). Both noctuid species have been observed immigrating overseas from southern and central China into South Korea and Japan (Wu et al., 2022; Otuka et al., 2023). The interpretation of the linear relationship between the δ^2^H values of waters and those of *M. separata* and *S. frugiperda* forewings requires clarification regarding the assimilation of hydrogen from waters in insect tissues for estimating natal origins of insect growth and ensuring the reproducibility of isotope analysis to investigate the source of long-distance migration.

We experimentally examined the effects of water on the δ^2^H values of maize leaves and *M. separata* and *S. frugiperda* forewings to reveal the linear relationship between the δ^2^H values of waters and those of maize leaves or insect forewings. For this purpose, we prepared five specific water samples for the cultivation of diet maize that imitated the gradient of hydrogen stable isotope ratios of environmental waters across the Japanese archipelago. Water sources and abiotic environmental factors facilitated the fluctuations of δ^2^H values of plants with seasonal changes in plant physiology. Our experimental design involved six cultivation periods with the rearing experiments. This enabled us to discuss the influence of plant physiology on the assimilation of hydrogen in insect tissues derived from water through feeding diets for the interpretation of the linear relationship between the δ^2^H values of the water samples and those of *M. separata* and *S. frugiperda* forewings.

## Materials and Methods

### Water sources

Bottled water from four commercial sources and tracer water (a mixture of deuterium and natural water) were used to cultivate maize for rearing *M. separata* and *S. frugiperda* larvae. Those four sources included three natural sources in Hokkaido, Yamagata, and Kumamoto prefectures, as well as one deep-sea water sample from the sea of Kochi Prefecture. The tracer water we prepared was 5 μL of deuterium water (99.9 atom % of deuterium, CAS no. 7789-20-0, Sigma Aldrich) per 1 L of commercial bottled water from Kumamoto. We sampled 6 mL of each bottle in each rearing experiment. All sampled waters were kept in a refrigerator at 4 °C until hydrogen isotope analysis.

### Cultivation of maize

In the rearing experiment, we used the maize cultivar LG3520 (Snow Brand Seed Co. LTD., Sapporo, Japan). Maize seeds in each water treatment were individually sown in a 16-hole cell box (#4401, Meiwa Co., LTD, Toyoake, Japan) with a commercial culture soil for vegetables (Protoleaf, Tokyo, Japan). The maize plants were watered daily. The seedlings were grown in a greenhouse at Kyushu Okinawa Agricultural Research Center (Koshi, Japan) and maintained under natural conditions from June to October 2020 (Table 1). There were a total of six cultivation periods, three for each species. For the *M. separata* rearing experiment, seeds were sown on 29 July and on 3 and 13 August 2020 (Table 1). For the rearing experiment of *S. furidperda*, seeds were sown on 14, 23, and 28 September 2020 (Table 1). Maize leaves were cut with scissors in a greenhouse, and the dust and soil were wiped away with 70% ethanol in a laboratory. All the maize leaves of each combination of cultivation period and water treatment were kept in a large plastic bag (Ziploc^®^, Asahi KASEI Co., Tokyo, Japan) in a refrigerator at 4 °C.

**Table 1.** Summary of each cultivation period of maize and the rearing periods of insects in 2020.

| Rearing Exp. | Developmental stage | Cultivation period | Sowing date | Supplying date | Length of cultivation period |
| --- | --- | --- | --- | --- | --- |
| *M. separata* | 1st – 3rd instars | Period 1 | 3 August | 11 or 18 August | 15 or 21 |
|  | 3rd – 5th instars | Period 2 | 29 July | 21 August | 23 |
|  | 6th instar | Period 3 | 13 August | 26 August | 13 |
| *M. separata* |  |  | Hatching date | Pupating date |  |
|  |  |  | 13 August | 8 September |  |
| *S. frugiperda* | 1st – 2nd instars | Period 4 | 23 September | 8 or 12 October | 15 or 19 |
|  | 3rd – 5th instars | Period 5 | 18 September | 16 October | 28 |
|  | 6th instar | Period 6 | 14 September | 22 October | 38 |
| *S. frugiperda* |  |  | Hatching date | Pupating date |  |
|  |  |  | 12 October | 27 October |  |

Length of cultivation days: this period refers to the days from the sowing date to the date when leaves were fed to *Mythimna separata* and *Spodoptera frugiperda* larvae. Egg masses of *M. separata* and *S. frugiperda* were set in cups on 11 August and 8 October 2020, respectively.

### Rearing conditions of *M. separata* and *S. furidperda*

We used successive strains of *M. separata* and *S. furidperda* in the rearing experiments. Approximately 100 larvae of each noctuid species were collected from maize pods at Kyushu Okinawa Agricultural Research Center in 2020. Fifty larvae were selected from each batch of 100 first instar larvae collected after hatching for each rearing experiment from each water source. These selected larvae were then reared in a clear plastic cup (Biocup, FG320TCL; Risu Pack Co., Gifu, Japan) until they reached the third instar stage. From the 4th to 6th instars, 20 larvae were selected and reared in a plastic case (150-1; Sanoya Co., LTD., Nagoya, Japan). Maize leaves were given daily during the larval stage in each water treatment. Pupae, until emergence, were individually transferred to a clear plastic cup (60TCL clean cup; Risu Pack Co., Gifu, Japan). The emerged adults were maintained without food for the third day and kept in a freezer at – 20°C for isotope analysis. All larvae were reared in an incubator (MIR-254-PJ; PHC Co., Tokyo, Japan), and the pupae were maintained in a growth chamber (PU-30; Koito Manufacturing Co., LTD., Tokyo, Japan) at 25°C under a 16-h light/8-h dark condition. At least 10 adult specimens were obtained in each water treatment in the rearing experiments.

### Isotope analysis

In preparation for isotope analysis, 1 mL of each water sample was transferred from a 6 mL glass vial to a 2 mL glass vial (e.g., Standard Vials, HIS3B18, Hitachi High-Tech Co., Tokyo, Japan) using a micropipette. The hydrogen stable isotope ratio of each bottled water was analyzed using cavity ring-down spectroscopy (L2140-*i*, Picarro, Inc., Santa Clara, CA, USA) at the Research Institute for Humanity and Nature (RIHN), Kyoto, Japan. The analytical precision values (1σ) of the L2140-*i* with liquid specifications were within 0.5‰ for δ^2^H values.

Maize leaves were cleaned with 70% ethanol and then were supplied to noctuid larvae. Scissors were used to remove the central vein from the maize leaf and to chop the maize leaf into small pieces. After that, pieces of maize leaves were placed in 1.5 ml tubes and dried at 60°C for 24 hours in a dry heat sterilizer (KM-300V, AS ONE, Co., Osaka, Japan). All dried samples were milled by using a bead shocker (Cell Destroyer PS1000, Bio Medical Science Co., Tokyo, Japan).

The forewings of *M. separata* and *S. frugiperda* were separated from the body with scissors, placed in 1.5 ml tubes, and dried at 100°C for 24 hours in the dry heat sterilizer. All maize leaves and forewing samples were prepared in a Kyushu Okinawa Agricultural Research Center laboratory.

In this study, the forewings of *M. separata* and *S. frugiperda* were not subjected to a rinse procedure using a methanol/chloroform solution (2:1, v/v) before for isotope analyses. Hairs, feathers, and tissue samples from wild animals are typically rinsed to remove soil and dust particles before analysis, as described in previous studies (e.g., Barbosa et al., 2009; Schnyder et al., 2006; Paritte & Kelly, 2009). In this study, however, all pupae were preserved without any residues in a clean plastic cup, and emerged adults were kept in a clean pack (Unipack®, Seisannipponsha LTD., Tokyo, Japan) at –20 °C until the stable isotope analysis.

A 0.25 mg portion of each sample was packed into a silver capsule (3.3 mm × 4.0 mm, Säntis Analytical AG, Landhausstrasse, Switzerland) by using an ultra-balance (XPR2U, Mettler Toledo, Greifensee, Switzerland). The δ^2^H values of the non-exchangeable hydrogen of maize leaves and forewings were determined at the Research Institute for Humanity and Nature (RIHN) with a comparative equilibration approach similar to that of Wassenaar and Hobson (2003). All samples and standard materials were placed in 96-well plastic ELISA plates. ELISA plates with cover were placed in a freeze-dryer (FDU-1200, EYELA, Tokyo Rikakikai Co., LTD., Tokyo, Japan) overnight to remove moisture from the sample surface, then loosely covered with the lid. Samples and keratin or cellulose standards were then allowed to equilibrate the ambient laboratory air moisture at room temperature for >96 hours. After the equilibration, ELISA plates were placed in a freeze-dryer overnight the day before the δ^2^H analysis, excluding the moisture on surface samples.

All measurements were performed by using a high-temperature conversion/elemental analyzer (TC/EA) (Thermo Fisher Scientific, Waltham, MA, USA) with a reduction unit equipped with a Costech Zero-Blank autosampler (Costech Analytical Technology, Inc., Valencia, CA, USA) and a Delta V advantage isotope-ratio mass spectrometer (IRMS) (Thermo Fisher Scientific). A chromium-filled reactor was employed to improve the reliability of δ^2^H analysis of nitrogen-bearing organic material (e.g., Coplen & Qi, 2012; Qi et al., 2016). A ceramic tube was filled from bottom to top with 0.5 cm of silver wool, 1 cm of quartz wool, 8 cm of glassy-carbon chips, 1 cm of quartz wool, 6 cm of chromium powder (500 μm, 99.0 % purity CAS-Nr.: 7740-47-3, Thermo Fisher Scientific), and 0.5 cm of nickel wool. The autosampler was purged and filled with dry helium gas to avoid the exchange of H_ex_ with ambient water. The reactor temperature was set at 1250 °C, and the GC temperature was maintained at 90 °C (Gehre et al., 2017). We cleaned the melted silver and sample residues every 200 to 250 samples from the tube. The measurement of δ^2^H values of maize leaves was corrected by the two-point calibration of USGS 54 (δ^2^H = −150.4‰, standard deviation: 1.5 ‰, n=30) and USGS55 (δ^2^H = −28.2‰, SD: 0.9‰, n=29) (Qi et al., 2016). The δ^2^H values of forewing samples were also calibrated by CBS (δ^2^H = −157.0‰, SD: 1.6‰, n=12) and KHS (δ^2^H = −35.3‰, SD: 1.4‰, n=12) (Soto et al., 2017).

Stable isotope ratios are expressed in delta notation in per mil (‰): δ^2^H (‰) = *R*_sample_/*R*_standard_ – 1, where *R* is the ratio of heavy to light isotopes (^2^H/^1^H). The isotope compositions of the present study are expressed relative to international standards, with hydrogen referenced to Vienna Standard Mean Ocean Water (VSMOW).

## Data analysis

All statistical analyses were performed using R ver 4.2.2. (R Core Team 2022). One-way ANOVA with Tukey’s multiple comparative method was used to analyze the effects of water on the δ^2^H values of maize leaves of each cultivation period as well as on *M. separata* and *S. frugiperda* forewings. Linear model (LM) analysis was conducted for relationships between the δ^2^H values of waters and maize leaves in each cultivation period or between the δ^2^H values of waters and those of *M. separata* and *S. frugiperda* forewings. Additionally, ordinal least squares regression using the LM was employed to analyze the relationships between the mean δ^2^H values of maize leaves in each cultivation period and those of forewings, accounting for the differing sample sizes of δ^2^H values between maize leaves and insect forewings.

## Results

### The δ^2^H values of bottled water

The δ^2^H values of five water samples ranged from –87.9 to 0.6‰ in the *M. separata* rearing experiment and from –87.7 to 2.8‰ in the *S. frugiperda* rearing experiment (Table 2).

**Table 2.**
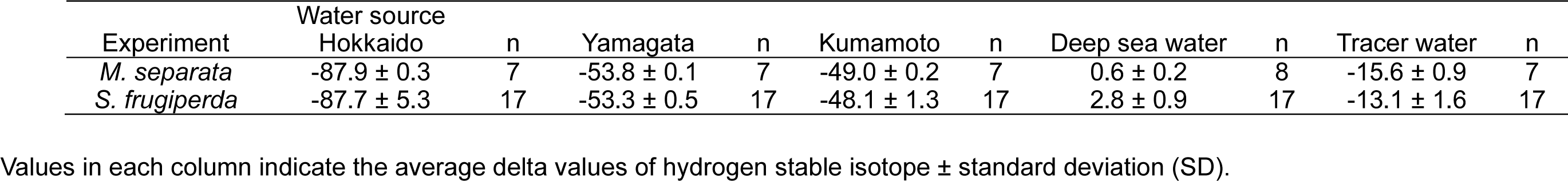
The δ^2^H values of water samples (‰) in the rearing experiments of *Mythimna separata* and *Spodoptera frugiperda* (Lepidoptera: Noctuidae).

### Effects of water on the δ^2^H values of maize leaves and insect forewings

The effects of water on the δ^2^H values of maize leaves and insect forewings were significant in both *M. separata* and *S. frugiperda* rearing experiments performed by one-way ANOVA with Tukey’s multiple comparative methods (Table 3). The δ^2^H values of maize leaves ranged from –64.4 to –38.1‰ in cultivation periods 1–3 and from –72.3 to –43.7‰ in cultivation periods 4–6 (Table 3). The δ^2^H values of the forewings ranged from –64.1 to –48.6‰ (*M. separata* experiment) and from –72.1 to –42.1‰ (*S. frugiperda* experiment) (Table 3).

**Table 3.** Results of one-way ANOVA with Tukey’s method to analyze the effect of water source on δ^2^H of maize leaves or forewings (‰) of *Mythimna separata* and *Spodoptera frugiperda* (Lepidoptera: Noctuidae) in each cultivation period.

|  |  | Water source |  |  |  |  |  |  |  |  |  |  |  |
| --- | --- | --- | --- | --- | --- | --- | --- | --- | --- | --- | --- | --- | --- |
|  |  | Cultivation period | n | Hokkaido |  | Yamagata |  | Kumamoto |  | Deep sea water |  | Tracer water |  |
| Experiment |  |  |  |  |  |  |  |  |  |  |  |  |  |
| *M. separata* | ML | Period 1 | 5 | -56.6 ± 5.3 | a | -50.2 ± 1.8 | b | -51.2 ± 2.7 | ab | -42.1 ± 2.9 | c | -40.4 ± 2.3 | c |
|  | ML | Period 2 | 4 | -63.7 ± 5.3 | a | -55.0 ± 3.2 | b | -53.4 ± 0.9 | b | -51.4 ± 2.5 | bc | -44.9 ± 1.4 | c |
|  | ML | Period 3 | 5 | -50.5 ± 3.8 | a | -44.6 ± 2.1 | b | -45.9 ± 2.1 | ab | -38.1 ± 2.9 | c | -36.0 ± 3.3 | c |
|  | Forewings |  | 10 | -64.1 ± 1.8 | a | -55.6 ± 2.0 | b | -55.5 ± 2.7 | b | -48.6 ± 3.6 | c | -49.8 ± 2.0 | c |
| *S. frugiperda* | ML | Period 4 | 5 | -63.9 ± 4.2 | a | -57.5 ± 1.9 | b | -55.5 ± 3.1 | b | -43.7 ± 4.2 | c | -48.0 ± 1.5 | c |
|  | ML | Period 5 | 5 | -72.3 ± 4.4 | a | -62.1 ± 3.6 | b | -59.5 ± 3.9 | bc | -50.3 ± 6.8 | d | -51.4 ± 3.5 | cd |
|  | ML | Period 6 | 5 | -70.6 ± 5.1 | a | -61.5 ± 3.2 | b | -56.3 ± 4.4 | bc | -44.5 ± 2.7 | d | -49.9 ± 2.5 | cd |
|  | Forewings |  | 10 | -72.1 ± 3.6 | a | -61.2 ± 3.4 | b | -56.0 ± 2.6 | b | -42.1 ± 2.8 | c | -48.9 ± 2.8 | d |

Values in each column indicate the average delta values of hydrogen stable isotope ± standard deviation (SD). N: the number of replications of forewings and maize leaf per water source. Abbreviation ML means maize leaves. The different letters in each row indicate $p < 0.05$ by Tukey's method in each sample.

### Relationships between δ^2^H values of water and those of maize leaves or insect forewings

All relationships between the δ^2^H values of water samples and those of maize leaves in each cultivation period, as well as between the δ^2^H values of water samples and those of insect forewings, showed significant linearities as determined by LM analyses (Table 4, Figures 1, 2). The slope value of the relationships between δ^2^H values of waters and those of organic compounds has been interpreted as a first-order approximation of the percentage contribution of environmental water to the tissue hydrogen content (see Ehleringer et al., 2008; Wang et al., 2009; Wolf et al., 2011). In cultivation periods 1–3, 16.2–18.6% of hydrogen in maize leaves was derived from water (Table 4). In cultivation periods 4‒6, 22.3–28.2% of hydrogen in maize leaves was derived from water (Table 4). The hydrogen content derived from water was calculated to be 17.4% for *M. separata* forewings and 32.7% for *S. frugiperda* forewings (Table 4).

**Figure 1.**
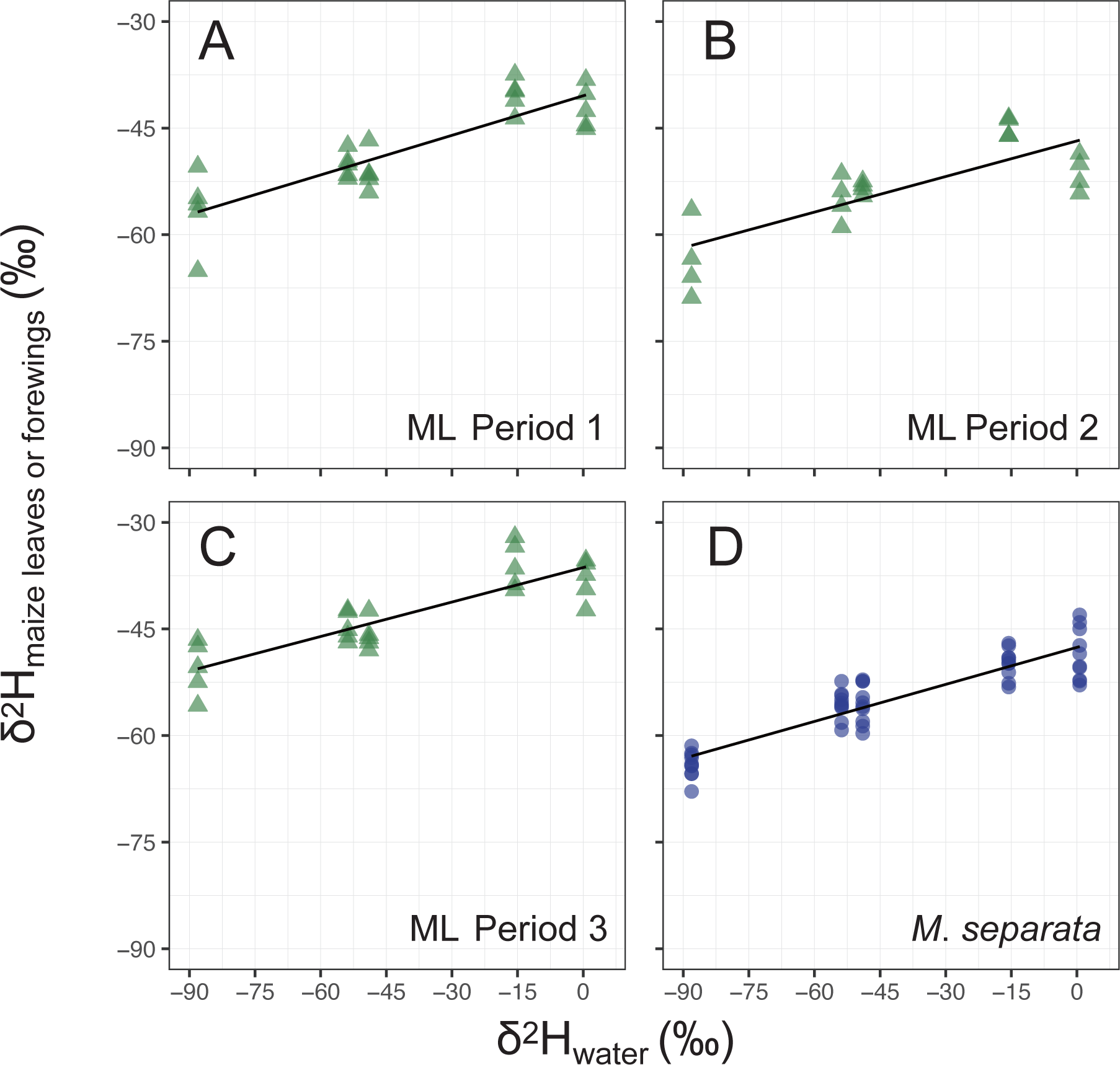
Relationship between the δ^2^H values of water sources and those of maize leaves in cultivation periods 1 (A), 2 (B), and 3 (C). Relationship between the δ^2^H values of water sources and those of forewings of *Mythimna separata* (D). Sample sizes of forewings and maize leaves are described in Table 3. Statistical results are summarized in Table 4.

**Figure 2.**
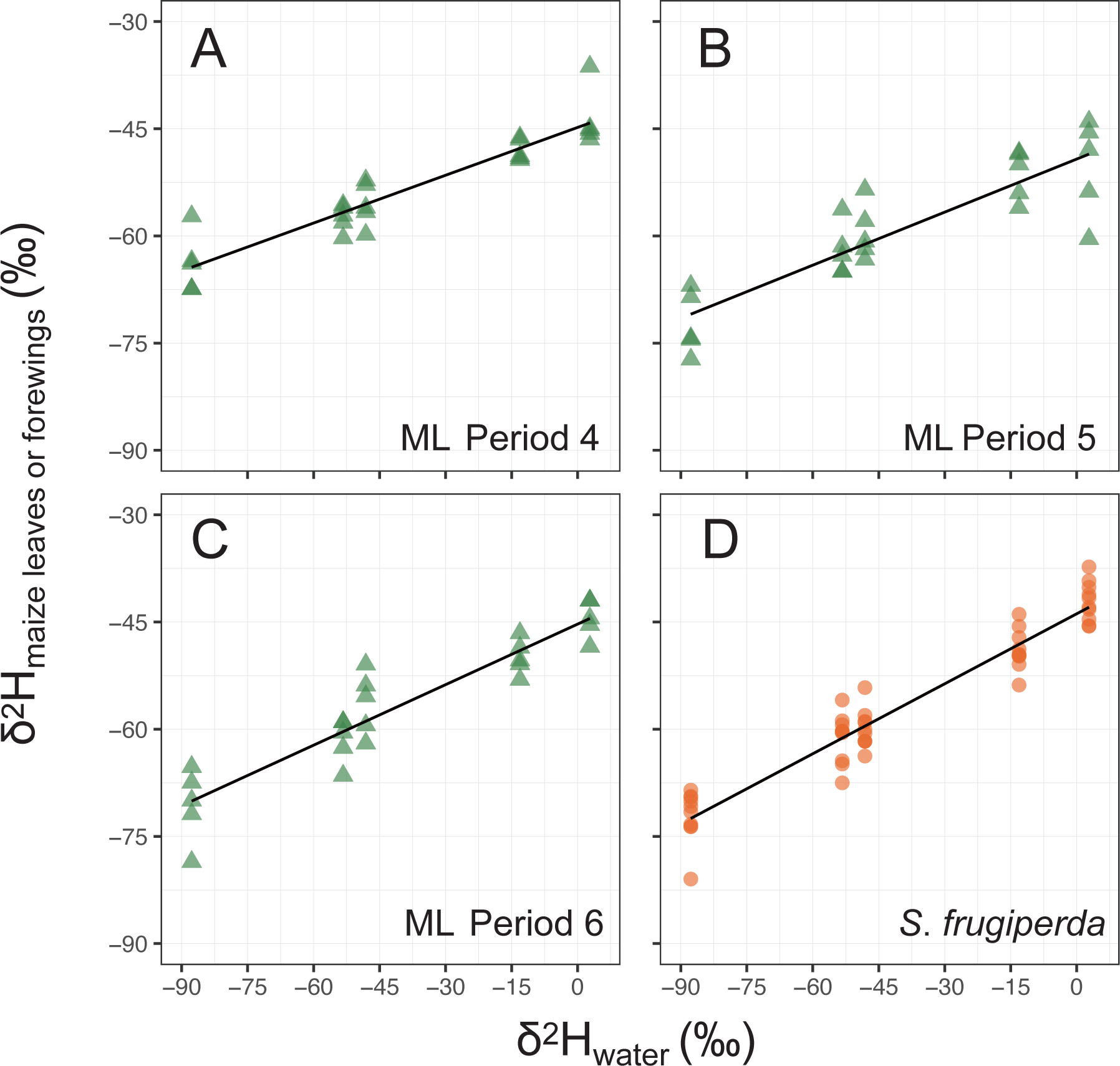
Relationship between the δ^2^H values of water sources and those of maize leaves in cultivation periods 4 (A), 5 (B), and 6 (C). Relationship between the δ^2^H values of water sources and those of forewings of *Spodoptera frugiperda* (D). Sample sizes of forewings and maize leaves are described in Table 3. Statistical results are summarized in Table 4.

**Table 4.** Summary of the linear regression results applied to the relationship between the δ^2^H values of water samples and those of forewings of *Mythimna separata* and *Spodoptera frugiperda* (Lepidoptera: Noctuidae) or maize leaves in each cultivation period.

| Experiment |  | Cultivation period | Slope value | Intercept | *P* | R<sup>2</sup> |
| --- | --- | --- | --- | --- | --- | --- |
| *M. separata* | ML | Period 1 | 0.186 | -40.449 | P<0.001 | 0.732 |
|  | ML | Period 2 | 0.167 | -46.803 | P<0.001 | 0.591 |
|  | ML | Period 3 | 0.162 | -36.363 | P<0.001 | 0.714 |
|  | Forewings |  | 0.174 | -47.588 | P<0.001 | 0.810 |
| *S. frugiperda* | ML | Period 4 | 0.223 | -44.819 | <0.001 | 0.854 |
|  | ML | Period 5 | 0.248 | -49.245 | <0.001 | 0.760 |
|  | ML | Period 6 | 0.282 | -45.297 | <0.001 | 0.858 |
|  | Forewings |  | 0.327 | -43.848 | <0.001 | 0.924 |

ML: maize leaves; OAW: *Mythimna separata*; FAW: *Spodoptera frugiperda*

### Relationships between δ^2^H values of maize leaves and those of *M. separata* and *S. frugiperda* forewings

All relationships between the mean δ^2^H values of maize leaves in each cultivation period and those of *M. separata* forewings showed significant linearities as determined by the LM with ordinal least squares (Table 5, Figure 3). The slope value of the linear regression between the mean δ^2^H values of maize leaves in cultivation period 3 and *M. separata* forewings was closest to 1.0 (slope value: 0.987) (Table 5). All relationships between the mean δ^2^H values of maize leaves in each cultivation period and those of *S. frugiperda* forewings showed significant linearities as determined by the LM with ordinal least squares (Table 5, Figure 3). The slope value of linear regression between the mean δ^2^H values of maize leaves in cultivation period 6 and *S. frugiperda* forewings was closest to 1.0 (slope value: 1.135) (Table 5).

**Figure 3.**
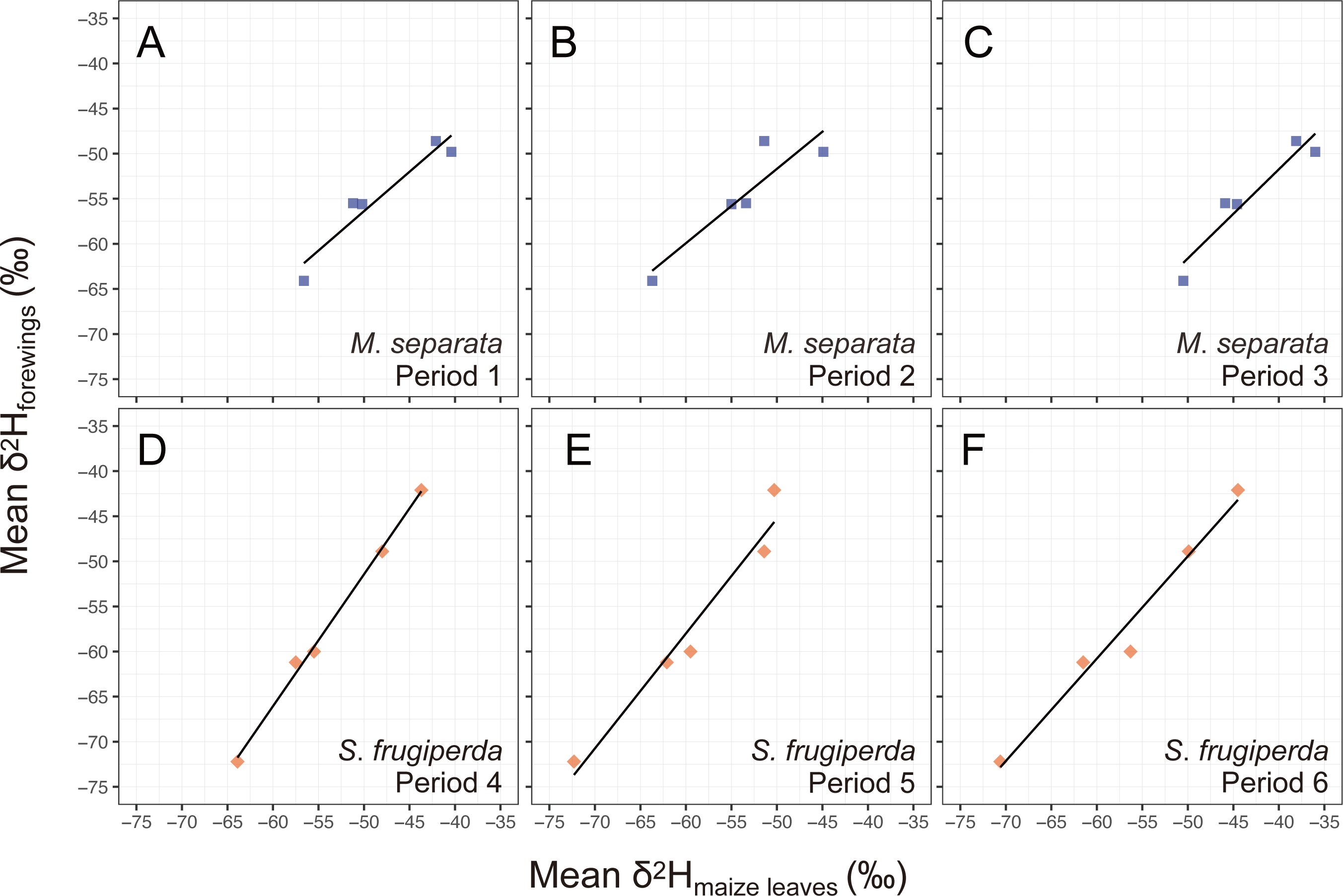
Relationship between the mean δ^2^H values of maize leaves in each cultivation period and those of forewings of *Mythimna separata* (A, B, C). Relationship between the mean δ^2^H values of maize leaves in each cultivation period and those of forewings of *Spodoptera frugiperda* (D, E, F). Statistical results are summarized in Table 5.

**Table 5.** Summary of the linear regression results applied to the relationship between the mean δ^2^H values of maize leaves in each cultivation period and those of forewings of *Mythimna separata* or *Spodoptera frugiperda*.

| Insect forewings | Cultivation period | Slope value | Intercept | *P* | *R*<sup>2</sup> |
| --- | --- | --- | --- | --- | --- |
| *M. separata* | Period 1 | 0.874 | -12.703 | <0.05 | 0.890 |
|  | Period 2 | 0.824 | -10.474 | <0.05 | 0.773 |
|  | Period 3 | 0.987 | -12.246 | <0.05 | 0.873 |
| *S. frugiperda* | Period 4 | 1.462 | 21.644 | <0.001 | 0.995 |
|  | Period 5 | 1.274 | 18.457 | <0.01 | 0.938 |
|  | Period 6 | 1.135 | 7.327 | <0.01 | 0.963 |

## Discussion

This study investigated the effects of different hydrogen isotope ratios in water on the δ^2^H values of maize leaves and the forewings of two noctuid species. The δ^2^H values of maize leaves and insect forewings ranged between –72.3 and –50.5‰ (Hokkaido), between –62.1 and –44.6‰ (Yamagata and Kumamoto), and between –51.4 and –38.1‰ (deep-sea and tracer water) (Tables 2, 3). The rearing experiments found linear correlations between the δ^2^H value of the water samples and those of maize leaves or *M. separata* and *S. frugiperda* forewings (Table 4, Figures 1, 2). The δ^2^H values of *M. separata* and *S. frugiperda* forewings were also linearly correlated with the δ^2^H values of maize leaves (Table 5, Figures 1, 2). These results are consistent with previous studies (Hobson et al., 1999b, 2018; Holder, 2012; Clem et al., 2022).

Our findings showed that 16.2–28.2% of hydrogen in maize leaves was derived from water through soils to root and plant bodies during cultivation periods. Temperature, humidity, precipitation, and other environmental variables directly affect water uptake by plants and facilitate the fluctuation of hydrogen isotope composition derived from water (Ehleringer & Dawson, 1992; Geris et al., 2017; Wang et al., 2017; Baan et al., 2023). Hydrogen in leaf tissues may be derived from photosynthetic carbon metabolism and ambient hydrogen through open stomas (e.g., Sanchez-Bragado et al., 2019). For example, 69% of the hydrogen in leaf cellulose in *Cryptomeria japonica* D. Don (Cupressaceae) was derived from ambient vapor, and 31% of the hydrogen originated from root water in the rainy season (Kagawa, 2022). The mean-percentage contribution of environmental water to maize leaves in cultivation periods 1–3 (17.6%) was lower than that in cultivation periods 4–6 (25.1%) (Table 4). The projection of the δ^2^H values of water to those of maize leaf differed among cultivation periods (Table 1). The fluctuation of δ^2^H values of *Juglans regia* L. (Juglandaceae) between 2017 and 2018 was affected by the amount of precipitation from July to August and the mean ambient air temperature (Krauẞ et al., 2020). The δ^2^H values of *Rubus idaeus* Linnaeus (Rosaceae), raspberry, and *Fragaria×ananassa* Duchesne (Rosaceae), strawberry, decreased with the increase in relative humidity in a laboratory chamber (Cueni et al., 2022). The δ^2^H values of cellulose sensitively responded to environmental differences during growing seasons (Baan et al., 2023). The present study suggested that changes in ambient temperature and seasonal humidity during cultivation periods may influence the exchange between hydrogen derived from water and ambient vapors in maize leaf tissues through the stomata during plant photosynthesis and respiration. Hydrogen derived from water in maize leaves may reflect physiological changes associated with seasonal variations during cultivation periods.

The LM analysis showed that *M. separata* and *S. frugiperda* assimilated 17.4% and 32.7%, respectively, of organic hydrogens derived from water through consumption of maize leaves (Table 3). Previous experiments revealed that the contribution of water to hydrogen in insect tissues was 40% in *Mythimna unipuncta* (Haworth) (Lepidoptera: Noctuidae) (Hobson et al., 2018) and 50% in *Danaus plexippus* (Linnaeus) (Lepidoptera: Nymphalidae), *Allograpta obliqua* (Say) (Diptera: Syrphidae), *Eupeodes americanus* (Wiedemann) (Syrphidae), and *Helicoverpa ameriga* (Hübner) (Noctuidae) (Hobson et al., 1999b; Holder, 2012; Clem et al., 2023). In the present study, the ratios of hydrogen isotopes derived from water in *M. separata* and *S. frugiperda* forewings were lower than those of other terrestrial insects (Hobson et al., 1999b; Holder, 2012; Clem et al., 2023). Hobson et al. (1999b) reported that the exchange between leaf water and ambient water vapor in the laboratory negatively affected the isotopic composition of hydrogen derived from water in *D. plexippus* wings. Another study indicated that changes in plant age and humidity influenced the fluctuation of δ^2^H values of *D. plexippus* wings from early to later eclosion dates (Lindroos et al., 2023). The exchange of water vapors in maize during cultivation periods may have affected the low ratios of hydrogen isotopes derived from water in *M. separata* and *S. frugiperda* forewings.

A previous study already indicated that there was almost a 1:1 relationship between δ^2^H values of *D. plexippus* wings and those of milkweed (Hobson et al., 1999b). In the present study, there was a near 1:1 relationship between the δ^2^H values of *M. separata* and those of maize leaves during cultivation period 3, and between the δ^2^H values of *S. frugiperda* and those of maize leaves during cultivation period 6 (Table 5, Figure 3). We provided these maize leaves planted during cultivation periods 3 and 6 to the final instar larvae of two noctuid moths (Table 1). We suggest that the hydrogen in *M. separata* and *S. frugiperda* forewings reflected the hydrogen in diet maize closely in the final instar larvae. The calibration of the natal origin of insects in field populations used the linear relationship between δ^2^H values of water samples and those of insect tissues (summarized in Hobson and Wassenaar 2018; Hobson et al., 2018). However, our study found that the host plant’s physiology affected the calibration equation of the insects. Therefore, careful consideration is required for interpreting the data of δ^2^H values of insect tissues, as the hydrogen isotope composition of insect tissues derived from waters may reflect the physiological status of the plant during the last instar larvae stage.

In conclusion, our experiments revealed the influence of host plant physiological changes on the interpretation of the linear relationship between the δ^2^H values of water and those of insect forewings. The contribution of hydrogen derived from water resources in forewings was lower than those of other terrestrial insects in previous studies. Water vapor in the atmosphere during plant photosynthesis and respiration might influence the exchange between hydrogen from water and ambient vapors in leaf tissues. As a result, the hydrogen isotope ratios derived from water samples in *M. separata* and *S. frugiperda* forewings were decreased. In addition, we found that the hydrogen isotope ratios of *M. separata* and *S. frugiperda* forewings most closely reflected those of maize leaves supplied during the final instar larval stage.

## Acknowledgment

TF and GA thank Dr. Naoki Kato for providing the maize seeds, S. Gyotoku for rearing the insects for the experiment, and Drs. Akira Otuka and Sachiyo Sanada-Morimura for directing grant projects and giving valuable comments on an early manuscript. This study was supported by a Joint Research Grant for the Environmental Isotope Study of Research Institute for Humanity and Nature to AO, by a Strategic International Collaborative Research project promoted by the Ministry of Agriculture, Forestry and Fisheries, Japan (JPJ008837) to SS-M, and by the research program on development of innovative technology grants (JPJ007097) from the Project of the Bio-oriented Technology Research Advancement Institution (BRAIN) to AO.

